# EnZymClass: Substrate specificity prediction tool of plant acyl-ACP thioesterases based on Ensemble Learning

**DOI:** 10.1101/2021.07.06.451235

**Authors:** Deepro Banerjee, Michael A. Jindra, Alec J. Linot, Brian F. Pfleger, Costas D. Maranas

**Affiliations:** The Bioinformatics and Genomics Program, Huck Institutes of the Life Sciences, The Pennsylvania State University, University Park, Pennsylvania, United States of America; Department of Chemical and Biological Engineering, University of Wisconsin-Madison, Madison, Wisconsin, United States of America; Department of Chemical Engineering, The Pennsylvania State University, University Park, Pennsylvania, United States of America

## Abstract

Classification of proteins into their respective functional categories remains a long-standing key challenge in computational biology. Machine Learning (ML) based discriminative algorithms have been used extensively to address this challenge; however, the presence of small-sized, noisy, unbalanced protein classification datasets where high sequence similarity does not always imply identical functional properties have prevented robust prediction performance. Herein we present a ML method, <u>En</u>semble method for en<u>Zym</u>e <u>Class</u>ification (EnZymClass), that is specifically designed to address these issues. EnZymClass makes use of 47 alignment-free feature extraction techniques as numerically encoded descriptors of protein sequences to construct a stacked ensemble classification scheme capable of categorizing proteins based on their functional attributes. We used EnZymClass to classify plant acyl-ACP thioesterases (TEs) into short, long and mixed free fatty acid substrate specificity categories. While general guidelines for inferring substrate specificity have been proposed before, prediction of chain-length preference from primary sequence has remained elusive. EnZymClass achieved high classification metric scores on the TE substrate specificity prediction task (average accuracy score of 0.8, average precision and recall scores of 0.87 and 0.89 respectively on medium-chain TE prediction) producing accuracy scores that are about twice as effective at avoiding misclassifications than existing similarity-based methods of substrate specificity prediction. By applying EnZymClass to a subset of TEs in the ThYme database, we identified two acyl-ACP TE, ClFatB3 and CwFatB2, with previously uncharacterized activity in *E. coli* fatty acid production hosts. We incorporated modifications into ClFatB3 established in prior TE engineering studies, resulting in a 4.2-fold overall improvement in observed C_10_ titers over the wildtype enzyme.

EnZymClass can be readily applied to other protein classification challenges and is available at: https://github.com/deeprob/ThioesteraseEnzymeSpecificity

**Author Summary:** The natural diversity of proteins has been harnessed to serve specialized applications in various fields, including medicine, renewable chemical production, and food and agriculture. Acquiring and characterizing new proteins to meet a given application, however, can be an expensive process, requiring selection from thousands to hundreds of thousands of candidates in a database and subsequent experimental screening. Using amino acid sequence to predict a protein’s function has been demonstrated to accelerate this process, however standard approaches require information on previously characterized proteins and their respective sequences. Obtaining the necessary amount of data to accurately infer sequence-function relationships can be prohibitive, especially with a low-throughput testing cycle. Here, we present EnZymClass, a model that is specifically designed to work with small to medium-sized protein sequence datasets and retain high prediction performance of function. We applied EnZymClass to predict the presence or absence of a desired function among acyl-ACP thioesterases, a key enzyme class used in the production of renewable oleochemicals in microbial hosts. By training EnZymClass on only 115 functionally characterized enzyme sequences, we were able to successfully detect two plant acyl-ACP thioesterases with the desired specialized function among 617 sequences in the ThYme database.

## Introduction

Machine Learning (ML) models are effective tools for narrowing the vast search space in complex biological problems. These techniques are especially effective in protein engineering and bioprospecting applications, as testing every possible residue substitution, or even every available homolog, is experimentally infeasible (1, 2). However, the efficacy of a ML model is dependent on the availability of an appropriately sized and balanced training dataset, with the requirement for known inputs scaling with the complexity and number of features needed to describe the data. Thus, a common barrier to utilizing data science for facilitation of experimental efforts is the compilation of a comprehensive dataset to train a predictive and accurate ML model. The development of a predictive model which can be trained with dataset sizes on the order of what is available to experimentalists, namely less than 1,000 training instances, would serve to facilitate a protein discovery and engineering pipeline for development of biocatalysts, biologic pharmaceuticals, and membrane transporters with a desired function. In this study, we focus on applying ML to the discovery of substrate-specific enzymes within the broader family of acyl- ACP thioesterases (TEs) using a training dataset of less than 120 characterized sequences in the academic and patent literature. We then use the predictions from EnZymClass to enhance the activity of the newly identified enzymes.

ML approaches which have been demonstrated to infer protein function and enzyme substrate specificity (3, 4) based on primary sequence fall under two categories, generative and discriminative. Generative approach methods build a model of the feature distribution for each protein category and assign a particular class or functional group to a candidate protein sequence by evaluating how well the sequence fits the model (5–12). Discriminative approaches on the other hand, focus on accurately learning the decision boundary between classes. Commonly used discriminative approaches rely on training Machine Learning (ML) classifiers such as Support Vector Machine (SVMs) or Neural Network (NNs) to learn discriminative rules from both positive (belonging to a particular protein class) and negative (not belonging to that protein class) set of protein sequences and apply the learnt rules to predict the class of any new protein sequence (13–15). While some of these studies have also incorporated structural information (16), ML algorithms have successfully identified pertinent sequence information to distinguish between highly similar proteins: guanylyl and adenylyl cyclases, lactate and malate dehydrogenases, and trypsins and chymotrypsins (17, 18).

Recent results suggest that discriminative approaches outperform generative approaches both in terms of accuracy and computational efficiency of solving the protein classification problem (19). SVM is among the most widely used discriminative learning algorithms for biological sequence classification (13–15,19–23) and has been experimentally proven to achieve up to 10% higher classification accuracy than generative approaches across a wide range of biologically relevant problems (20). While effective, SVM classifiers are highly influenced by the feature extraction technique employed to encode the protein sequences (24). Proper selection of a feature extraction method is integral to the model performance, especially when available training data is limited.

Feature extraction of protein sequences generates a discrete numerical representation of a protein to create feature vectors that are correlated with the attribute of the protein that will be predicted by the model. In order to train an SVM, several feature extraction techniques for protein sequences have been implemented. These techniques can be divided into four categories: kernel-based methods, physicochemical encoding of protein sequences, N-gram representations, and Position Specific Scoring Matrices derived methods. Kernel-based methods such as the spectrum kernel (14) or its more generalized form, the mismatch kernel (19), both introduced by Leslie et. al., calculate the number of times all possible *k*-mers or *k*-length contiguous subsequences occur in a given protein sequence. Apart from kernel-based methods, another class of feature representation techniques extract structural and physicochemical properties embedded in the protein sequence and convert it into a numerical vector. Amino Acid Composition (AAC) descriptor developed by Nakashima et. al. (25) and Composition-Transition-Distribution (CTD) descriptor developed by Dubchak et. al. fall under the second class of protein sequence encoders (13). The third category, N-gram representation, is derived from Language models (26) which assumes that an amino acid is analogous to a word in a sentence. The n-gram model assigns probabilities to *n* contiguous sequences of amino acids based on their occurrence in a set of protein sequences and uses the probabilities to create a numerical representation of a protein sequence. Features have also been derived from PSSM profiles, which incorporate evolutionary information about a protein sequence by deriving position specific substitution scores from multiple sequence alignment of other highly similar protein sequences present in a database (27). While selection of the most informative feature extraction technique can improve performance of a classifier, performance tends to be problem specific without being generalizable across the entire protein classification domain. Nanni *et al.* recognized this limitation and suggested the use of an ensemble of classifiers, each trained on separate feature descriptors of protein sequences, to attain consistently superior performance across the entire protein classification domain compared to any individual feature extraction technique (28).

Several studies have shown that ensemble methods performed better than any individual classification method especially in problems relevant to the protein classification domain (29–32). Camoglu *et al.* used a decision tree based ensemble classifier to classify proteins in the Structural Classification of Proteins database and showed how it is possible to attain much lower error rates using the ensemble classifier than any individual method (30). While solving the protein fold classification problem, Tan *et al.* illustrated the advantage of using ensemble classifiers on unbalanced datasets (32). Similarly, Caragea *et al.* trained an ensemble of SVM classifiers to predict glycosylation sites in amino acid residues and found that an ensemble of SVMs outperformed an individual SVM trained on imbalanced data (33). We thus set out to demonstrate the utility of an ensemble of discriminative ML classification algorithms to broadly predict the presence or absence of a useful protein characteristic from primary sequence information. We selected the detection of medium-chain-length specificity of acyl-ACP TEs as our classification problem due to 1) the relatively small number of experimentally characterized medium chain acyl-ACP TE sequences, 2) the importance of the TE in medium-chain fatty acid synthesis, and 3) the absence of any existing computational tools for this task.

Medium-chain oleochemicals, defined as eight to twelve-carbon free fatty acids and derivatives, are target molecules for synthetic biologists due to limited or challenged accessibility from conventional agricultural or petrochemical routes (34–38). While these chain lengths have traditionally been sourced from tropical crops, such as palm, palm kernel, and coconut, the eight, ten, and twelve-carbon products are not major constituents of the oil (39). Furthermore, the displacement of rainforest habitat due to the cultivation of the oil palm has been identified as the single largest impact on decreasing biodiversity observed in the Southeast Asian jungle ecosystem (40). Processes have been established to create higher value oleochemical derivatives, such as fatty alcohols, directly from petrochemical building blocks. However, these processes yield a distribution of alcohols, and thus do not provide a highly selective route to the medium-chain products (41).

Using the tools of synthetic biology, fatty acid and fatty alcohol distributions with over 90% of the product belonging to the C_8_ species have been attained (42, 43). This has been achieved via rewiring of the fatty acid biosynthesis pathway in *E. coli*, namely by the incorporation of an engineered eight-carbon specific acyl-ACP TE from *Cuphea palustris*. Indeed, the expression of various acyl-ACP TEs, either homologs from nature or variants thereof, has enabled control over the chain-length distribution in *E. coli* production systems (42–47) **(Figure 1)**. Of these studies, acyl-ACP TEs from select plant species have been shown to have greater native specificity toward the medium-chain substrates when compared to bacterial homologs (38,45,48,49). The *Cuphea* genus, a flowering shrub whose seed oils commonly exhibit over 70% of medium-chain triglycerides, has served as source of these TE genes (50). Indeed, many homologs from these shrubs have been patented for expression in bacteria and algae due to their utility in redirecting the flux of carbon in fatty acid biosynthesis to medium-chain fatty acid derivatives (51, 52). While substrate specificity is generally understood to be influenced by the binding pocket and the acyl carrier protein landing pad, prediction of acyl-ACP TE substrate chain-length preference *a priori* remains an unsolved challenge. In bioprospecting efforts, it is typical that multiple TE genes are isolated in cDNA libraries prepared from plant tissue extracts. Thus, the identification of the gene responsible for the medium-chain activity requires further characterization such as *in vivo* production experiments in a genetically tractable host or *in vitro* activity assays (53). Therefore, discovery of novel acyl-ACP TEs is desirable to both navigate the current intellectual property landscape in oleochemical biosynthesis and to elucidate the mechanisms of substrate specificity among this enzyme class.

**Figure 1:**
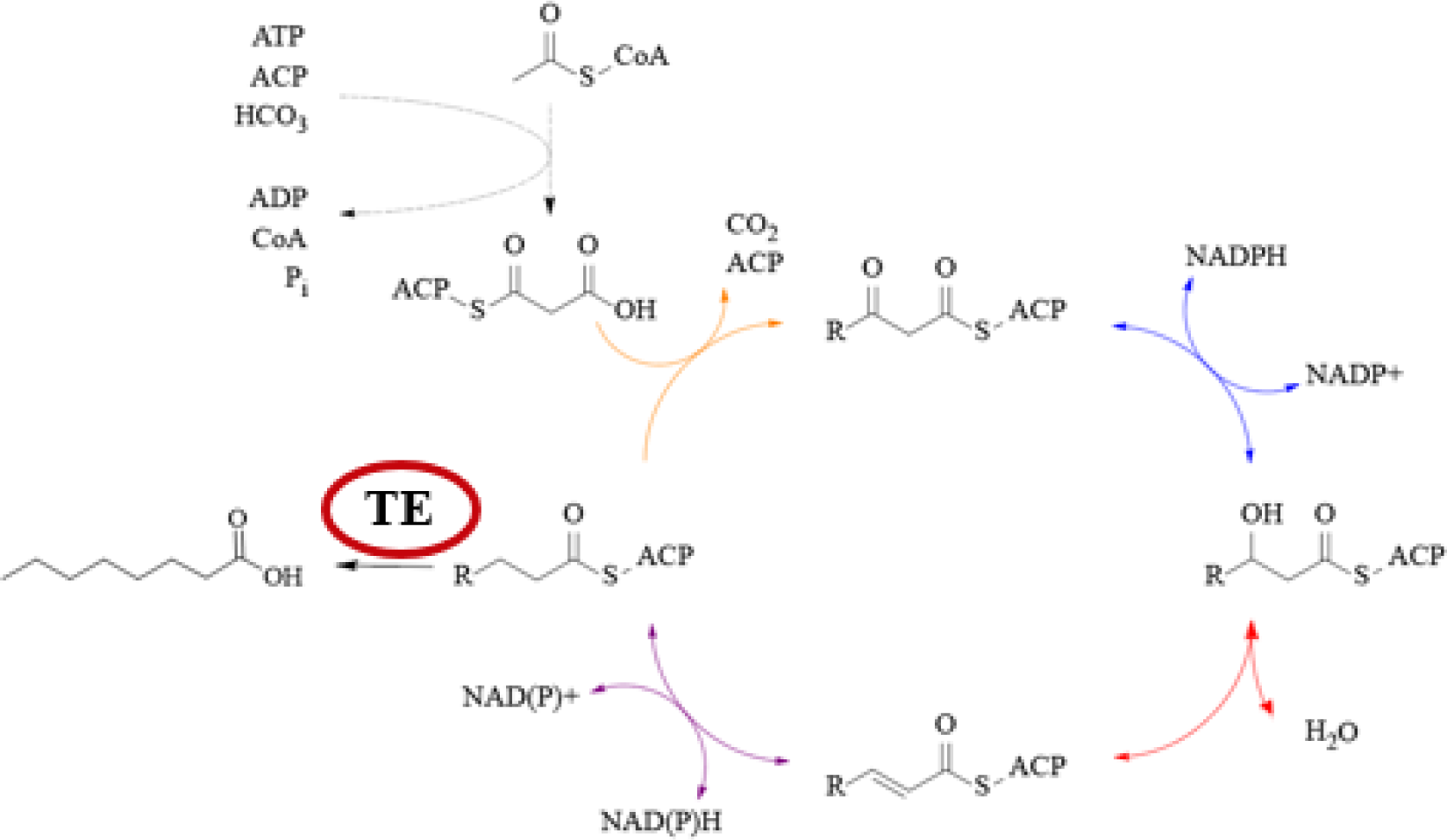
The acyl-ACP TE plays a key role in fatty acid biosynthesis in E. *coli*. By intercepting the growing acyl-ACP chains, the TE hydrolyzes the acyl chain from the ACP and redirects flux to the free fatty acid pool. These free fatty acids can be further derivatized *in vivo* or *ex vivo*.

To bridge the sequence-to-function mapping gap, the ThYme database was compiled to organize putative and characterized thioester-active enzymes into various families based on their primary sequence information (54). As a follow-up study, Jing *et al.* characterized the *in vivo* activity of 24 TEs from the TE14 family in the ThYme database and showed that phylogenetic and sequence identity analysis alone were not sufficient to distinguish plant TEs substrate specificity (46).

Herein we put forth a machine learning based discriminatory approach, termed EnZymClass (**En**semble method en**Zym**e for **Class**ification), to predict substrate specificity from primary sequence encoded features for uncharacterized TEs. EnZymClass is a stacked ensemble classifier comprised of 47 base learners that are trained using 47 different feature extraction techniques (listed in **Table 2**) which numerically encode enzyme primary sequences. It relies on a meta learner which automatically selects the top five base learners and combines their output by applying a hard majority voting criterion to predict the substrate specificity class of a TE. Classification models that perform well across multiple protein classification datasets and application areas have been previously implemented (28). However, methods that explicitly deal with the challenges of high-dimensionality, small size, dataset imbalance, and sequence similarity, all commonly encountered in protein classification tasks, remain scarce. The framework developed in this work is carefully designed to address these issues at every stage of its pipeline. EnZymClass achieved a mean validation accuracy of 0.8, mean precision and recall scores of 0.87 and 0.89 on medium chain TE category prediction, the primary product of interest in this study. There presently exists no computational method capable of performing accurate large-scale categorization of plant acyl TEs into their substrate specificity groups. We overcame this limitation by using the developed ensemble method to identify two medium-chain acyl-ACP TEs, ClFatB3 and CwFatB2, among a set of 617 TEs catalogued in the ThYme database. We further modified the gene sequence of ClFatB3 to improve titer of ten-carbon free fatty acids by 4.2-fold over wildtype (WT) when expressed in an *E. coli* production host. This study provides an exemplar of how even limited datasets can be leveraged through ML to support bioprospecting efforts and to provide a suitable starting template enzyme for protein engineering experiments. EnZymClass can be accessed at: https://github.com/deeprob/ThioesteraseEnzymeSpecificity

## Results

### EnZymClass classification of TEs

We evaluated the performance of EnZymClass on TE substrate specificity prediction task using a rigorous validation scheme (discussed in Model evaluation) to fairly assess model generalizability, robustness, and reproducibility of results. EnZymClass achieved a mean validation accuracy of 0.8 across 10,000 simulations using different training and validation sets. Although, the worst-case accuracy across simulations was 0.52, the standard deviation of accuracy distribution was only 0.06 which implies robustness to training set selection. The mean precision score of EnZymClass across the simulations for the medium-chain TEs, the product of interest, was 0.87 while it was 0.78 for the long-chain specific TE and 0.54 for the TEs of mixed specificity. The mean recall scores of the ensemble for the medium-chain, long-chain and mixed specificity TEs were 0.89, 0.92, and 0.28, respectively. The mean Matthew’s Correlation Coefficient (MCC) validation score of EnZymClass was 0.68. The precision (on the medium-chain category), recall (on the medium- chain category), accuracy, and MCC score distributions of EnZymClass are shown in **Figure 2**.

**Figure 2:**
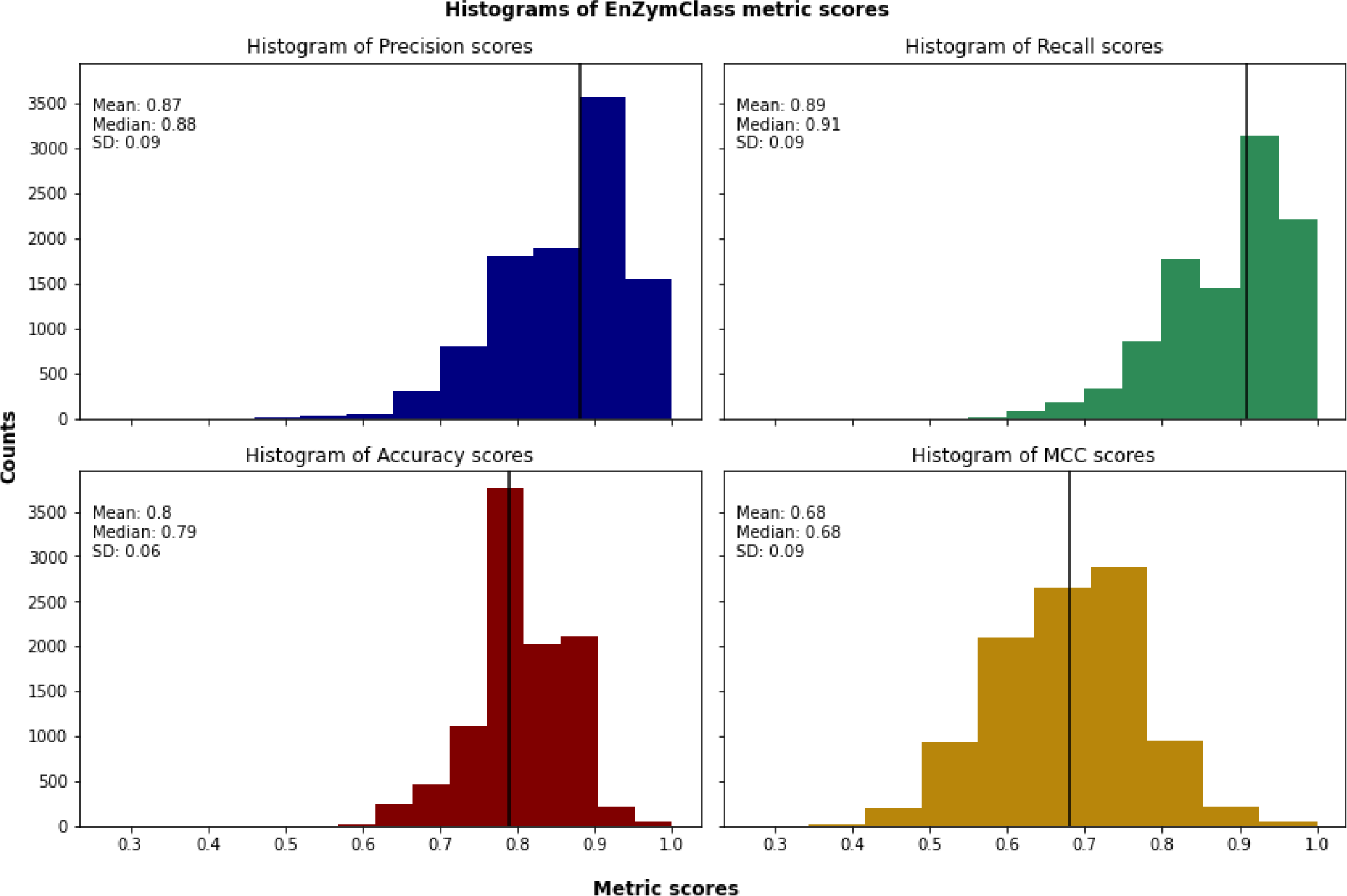
Precision (on medium-chain TEs), Recall (on medium-chain TEs), Accuracy, and MCC score distribution of EnZymClass. The black vertical line denotes the median score of the specific metric. The mean, median, and worst precision scores are 0.87, 0.88, and 0.40, respectively. The mean, median, and worst recall scores are 0.89, 0.91, and 0.50, respectively. The mean, median, and worst accuracy scores are 0.8, 0.79 and 0.52, respectively. The mean, median, and worst MCC scores are 0.68, 0.68 and 0.27, respectively.

Among the three categories of TEs, the mixed specificity class is the worst predicted with much lower precision and recall scores compared to the medium-chain and long-chain categories. This may be attributed to the fact that the mixed specificity category of TEs represents only 17% of all the characterized TEs in our dataset. Although EnZymClass offers a mechanism to deal with class imbalance by allowing the user to provide label priority weights as base learner input that in turn manipulates the learner to penalize a label misclassification based on those weights, we opted against assigning such weights to the minority class. Prioritizing correct classification of the mixed specificity class of TEs would have de-emphasized classification of medium-chain TEs thus impacting its accurate characterization. The base learners in the ensemble were chosen as SVM classifiers after analyzing the performance of SVMs against two learning algorithms: GBCs and NNs. GBCs and NNs were both outperformed by SVM in terms of accuracy on separate held-out validation sets (**Table S2**). SVMs have higher generalizability and can handle high dimensional datasets (55) which may have resulted in its superior performance on our relatively small dataset. In comparison to existing studies that aim to characterize substrate specificity of TE sequences, EnZymClass can perform at a much larger scale enabling identification of TE chain-length specificity across large databases such as ThYme (54). Existing efforts to characterize TEs mostly rely on experimental approaches which limit their ability to test at most ten to thirty TEs at a time (56–59). Although phylogenetic analysis and sequence alignment-based approach have been used before (46, 60), they were mostly restricted to a few specific classes of TEs. Nevertheless, previous studies have demonstrated that phylogenetic analysis or sequence similarity-based approaches do not always correlate with TE substrate specificity (46, 59). At present, there is no available computational platform that can match the TE substrate specificity characterization of EnZymClass.

### Comparison with individual base learners in EnZymClass

The primary purpose of using an ensemble framework in EnZymClass was to decrease the amount of variance in model prediction resulting from a small training set. In addition, ensemble methods can improve model performance and increase model robustness to a limited and unbalanced training set (30–32). Our results indicate that the ensemble model delivered on all three fronts, performing better than any individual base model trained on a specific feature extraction technique. It achieved a mean accuracy of 0.8, mean precision (on medium-chain TEs) of 0.87, and mean recall score (on medium-chain TEs) of 0.89 on varying validation datasets across 10,000 simulations of our study. In comparison, the three best performing individual base models trained on Spectrum Kernel, Gappy Kernel and CKSAAP feature extraction techniques using the SVM learning algorithm achieved mean accuracy scores of 0.77, mean precision (on medium-chain TEs) scores of 0.87 and mean recall (on medium-chain TEs) of 0.85, 0.86, and 0.86, respectively on the same validation datasets. A detailed analysis of the 47 base models used in EnZymClass, each trained on a unique feature extraction technique is provided in **Table S1**. EnZymClass also showed an increased robustness to training set significantly improving the worst-case accuracy score achieved over the 10,000 varying validation datasets from 0.48 (highest worst-case accuracy obtained by any base model) to 0.52. Moreover, the standard deviation of the distribution of mean prediction accuracy also decreased from 0.07 (least mean accuracy score variance achieved by any base model) to 0.06. Similar improvements were found for the recall score metric on medium-chain TEs as well. A comparison of the mean, minimum, and standard deviation scores of the three classification metrics under consideration (i.e., mean accuracy, precision and recall score on medium chain TEs) between EnZymClass and the five top performing base models (based on validation data scores) is tabulated in **Table 1**.

**Table 1:** EnZymClass performs better than any individual base model on varying validation datasets in terms of prediction accuracy and robustness to training set. We illustrate that phenomenon by comparing ensemble model results with the results of five best performing base models (judging by their performance on validation datasets). Three popular classification metrics, accuracy, precision, and recall, were used to compare the models. The mean, minimum, and standard deviation of the three metrics achieved by EnZymClass and the five best performing base learners on varying validation datasets are displayed here.

| Model | Mean | Min | Std | Mean | Min | Std | Mean | Min | Std |
| --- | --- | --- | --- | --- | --- | --- | --- | --- | --- |
| Name | Precision | Precision | Precision | Recall | Recall | Recall | Accuracy | Accuracy | Accuracy |
| EnZymClass | 0.87 | 0.40 | 0.09 | 0.89 | 0.5 | 0.09 | 0.8 | 0.52 | 0.06 |
| Spectrum<br>Kernel | 0.87 | 0.44 | 0.09 | 0.85 | 0.36 | 0.10 | 0.77 | 0.45 | 0.07 |
| Gappy<br>Kernel | 0.87 | 0.44 | 0.09 | 0.86 | 0.38 | 0.10 | 0.77 | 0.48 | 0.07 |
| CKSAAP | 0.87 | 0.38 | 0.09 | 0.86 | 0.35 | 0.10 | 0.77 | 0.45 | 0.07 |
| KSCTriad | 0.86 | 0.33 | 0.09 | 0.85 | 0.33 | 0.10 | 0.76 | 0.41 | 0.07 |
| Moran | 0.87 | 0.33 | 0.10 | 0.85 | 0.38 | 0.10 | 0.76 | 0.45 | 0.08 |

Among the 47 base models trained in EnZymClass, the Spectrum Kernel descriptor trained model produced the best accuracy scores, followed by Gappy Kernel and Composition of *k*-spaced amino acid pairs (CKSAAP). Kernel-based methods primarily calculate the counts of all possible *k*-length contiguous subsequences present in a protein sequence. CKSAAP is a general form of the Dipeptide composition (DPC) descriptor but differ by calculating the composition of amino acid pairs, separated by at most a distance of *k*. DPC, however, only calculates the composition of neighboring amino acid pairs. Interestingly, DPC is also among the top-performing descriptors in terms of average accuracy scores (ranked sixth), further emphasizing the efficacy of dipeptide composition as predictors of substrate specificity in TEs. A list of the 47 feature encoding techniques trained base models ranked by their mean accuracy scores on TE substrate specificity prediction task is given in **Table S1**.

### Comparison with similarity-based classification method

We further benchmarked EnZymClass by comparing it to an existing sequence similarity-based classification method. Sequence similarity-based methods define a distance function to measure the similarity between a pair of sequences (23). Here, we calculated the BLASTP (61) identity scores between a pair of subject and query sequence to use as the distance function. Henceforth, we trained a *k*-Nearest Neighbors classifier with *k* set as three on a subset of TE sequences and predicted the substrate specificity on the remaining validation set. This process was repeated 10,000 times by varying the training and validation sets using different random seeds similar to EnZymClass evaluation scheme (as discussed in the Model evaluation section). The sequence similarity-based model achieved a mean accuracy score of 0.70, mean precision score (on the medium-chain TEs) of 0.79, and mean recall score (on the medium-chain TEs) of 0.79. In comparison, EnZymClass attained a mean accuracy score of 0.80, mean precision score (on the medium-chain TEs) of 0.87, and mean recall score (on the medium-chain TEs) of 0.89, producing significantly better results. The distribution of the precision, recall, and accuracy scores for the similarity-based classification model along with the box plots comparing those distributions with the similar validation score distributions achieved by EnZymClass is shown in **Figure 3**.

**Figure 3:**
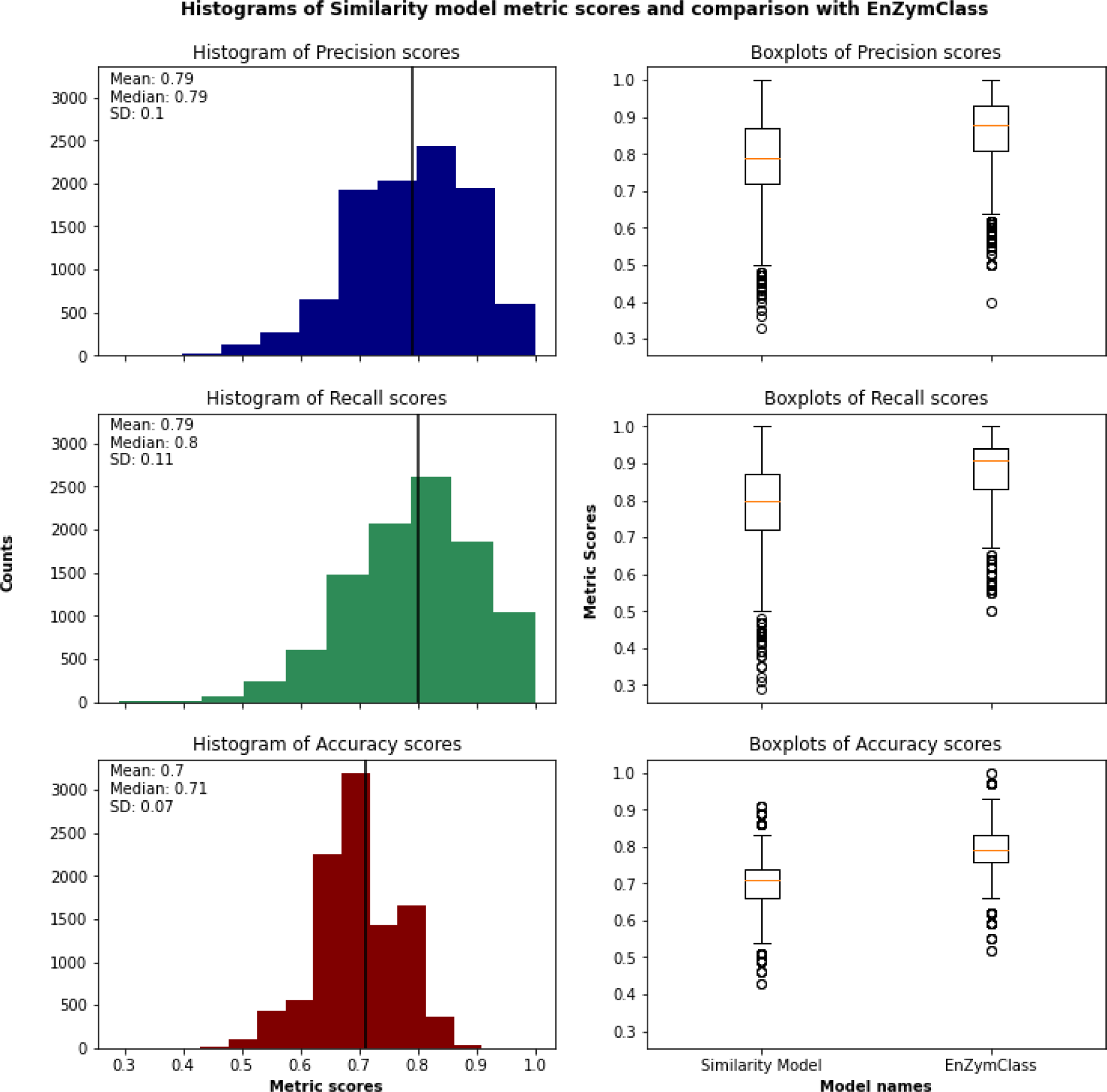
Precision (on medium-chain TEs), Recall (on medium-chain TEs), and Accuracy score distribution of the similarity model along with the comparisons of those distributions with EnZymClass through box plots is depicted. The mean precision, recall, and accuracy scores are 0.79, 0.79, and 0.7, respectively. The box plot comparisons between Similarity model and EnZymClass performance metrics show the EnZymClass metric distributions are skewed towards higher scores across all metrics, thus asserting the superiority of EnZymClass over similarity-based model.

Previous studies indicated that sequence similarity is not the best predictor of TE substrate specificity since similar sequences may have different substrate specificity (46, 59). For example, the TEs from *Cuphea viscosissima* share more than 70% sequence identity but display varying substrate specificities whereas TEs from *Cocos nucifera* and *Sorghum bicolor* have similar substrate specificity although they share less than 40% sequence similarity. Feature extraction techniques from protein sequences used in EnZymClass, not only encode similarity between sequences but also extract physicochemical, contextual, and evolutionary information from protein sequences. The sequence similarity-based model results delivered further evidence in support of the “similarity does not infer specificity” argument while justifying the use of protein sequence feature extraction techniques as better predictors of TE substrate specificity.

### Identification of two uncharacterized medium-chain TEs from the Cuphea genus

EnZymClass predicted three enzyme sequences to encode medium-chain length specific TEs from the TE14 Family in the ThYme database (54). The TE14 Family contained about 2,500 sequences of prokaryotic and eukaryotic sequences, however, since the training set was comprised of solely eukaryotic sequences from the plant kingdom, the search was restricted to the subset of 617 sequences with eukaryotic origin. The *in vivo* performance of the predicted TEs when expressed in the *E. coli* RL08*ara* strain is summarized in **Figure 4a** and **Table S4**. The three TEs, ClFatB3, ClFatB3-2, and CwFatB2 are from various *Cuphea* species: the former two from *Cuphea lanceolata* and the latter from *Cuphea wrightii*. ClFatB3 and CwFatB2 showed medium-chain length activity. ClFatB3-2 yielded a titer and distribution similar to the catalytically inactivated control, BTE-H204A, and thus produced no medium-chain free fatty acids. ClFatB3-2 differed from ClFatB3 by one amino acid in the binding pocket: position 135 in ClFatB3 is a serine while position 135 in ClFatB3-2 is a proline.

**Figure 4:**
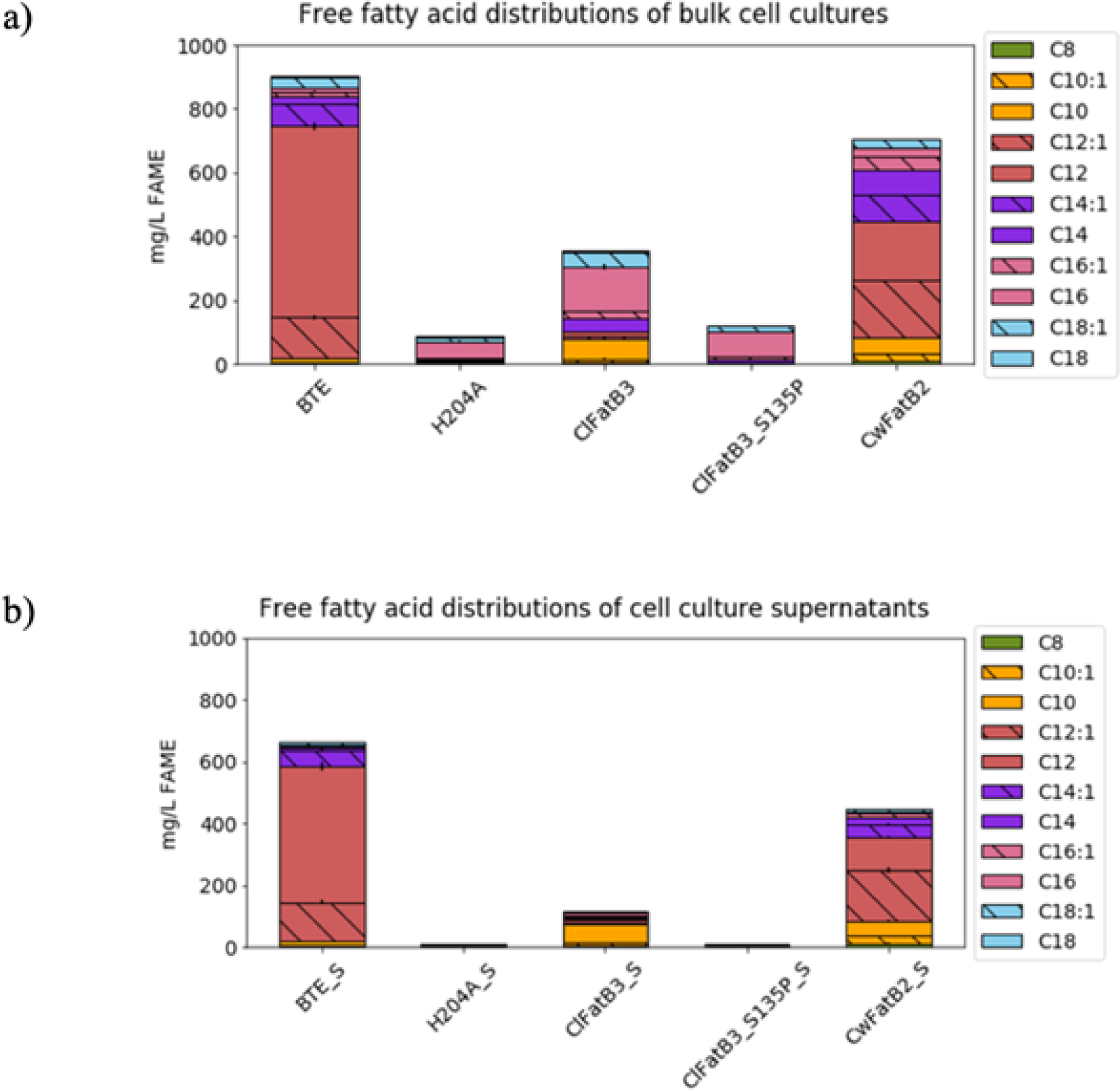
The effect of TE homolog on free fatty acid distribution in a) bulk cultures and b) supernatants of RL08*ara E. coli* cells. The distribution from cells expressing the California bay laurel TE (BTE) and a catalytically inactive BTE variant (H204A) are shown as a positive and negative control, respectively. Bar height represents the average titer obtained from biological triplicates, and error bars represent the standard error of the mean.

The free fatty acid distributions from the *in vivo* cultures contained a significant proportion of hexadecanoic and hexadecenoic acid. This was particularly apparent in the cultures expressing the ClFatB3 homolog. We reasoned that these species were cell membrane constituents which resulted from derivatization of phospholipids rather than free fatty acids. To validate this assumption, the cultures were centrifuged, and the supernatant was collected to be analyzed for free fatty acid content. The results from this processing step shown in **Figure 4b** and **Table S5** confirmed that ClFatB3 and CwFatB2 are primarily medium-chain specific, demonstrating the efficacy of EnZymClass in identifying substrate preference. While ClFatB3-2 did not exhibit medium-chain specificity, the analysis of the supernatant suggested this was due to enzyme inactivity. Since ClFatB3-2 only differed from ClFatB3 by a single residue, EnZymClass also classified ClFatB3 as a medium-chain specific TE. This instance illustrates the limitations of EnZymClass applied in this study, as it was tuned to only classify if a TE exhibits medium-chain, long-chain or mixed substrate specificity. The addition of another classifier for predicting inactive TE could be in principle possible provided an inactive TEs training dataset.

### Improving activity of medium-chain acyl-ACP TE identified from ML predictions

We sought to improve the activity of the ClFatB3 and CwFatB2 enzymes by incorporating mutations previously demonstrated to be effective for enhancing free fatty acid production. Hernández-Lozada *et al.* leveraged an auxotrophic *E. coli* strain to screen for mutations productive toward enhancing activity of an octanoyl-ACP TE from *Cuphea palustris*, CupTE (42). This resulted in the isolation of a variant TE with a fifteenfold improvement in k*_cat_* for the native substrate over the WT. The three mutations which conferred the increased activity included a truncation, a mutation near the N-terminus, and a mutation in the binding pocket (ΔA_54_, N28S, and I65M, respectively). Since acyl-ACP TE truncations have been shown to be effective in varying activity (62), the ΔA_54_ truncation was first incorporated into all three TE sequences identified from the model predictions as well as to the positive BTE control sequence. The effect of this truncation on free fatty acid distribution is shown in **Figure 5** and in **Table S4**. The Clustal Omega multiple sequence alignment used to locate the truncation site is shown in **Figure S1** (63).

**Figure 5:**
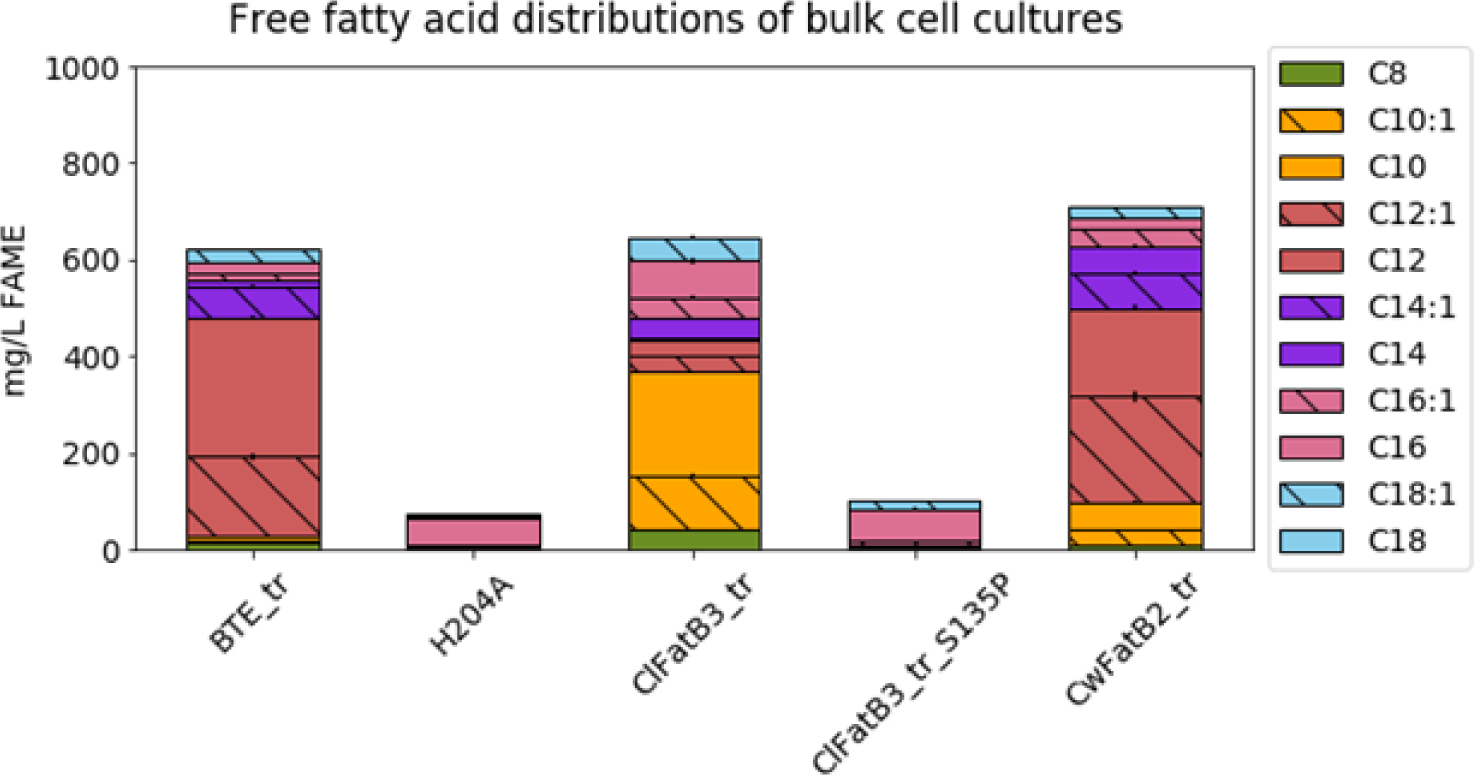
The effect of TE homolog on free fatty acid distribution in bulk cultures of RL08*ara E. coli* cells. Each TE was truncated at the location where the N-terminus aligned with the ΔA_54_ truncation in CupTE. The distribution from cells expressing the California bay laurel TE (BTE) and a catalytically inactive BTE variant (H204A) are shown as a positive and negative control, respectively. Bar height represents the average titer obtained from biological triplicates, and error bars represent the standard error of the mean.

The effect of the truncation was most beneficial to the ClFatB3 enzyme, resulting in a 3.3-fold increase in combined decanoic and decenoic acid titers. The CwFatB2_tr variant displayed a 5% increase of dodecanoic acid in the free fatty acid distribution when compared to the non-truncated sequence but did not exhibit an improvement in overall titer. The ClFatB3-2 TE sequence was also truncated to determine if its inactivity could be overcome by the N-terminal modification. However, no change was observed.

### Incorporation of activity-enhancing mutations into ClFatB3

We sought to further enhance the activity of ClFatB3 for two reasons: 1) the N-terminal truncation resulted in a 3.3-fold improvement in C_10_ free fatty acid titer, and 2) no decanoyl-ACP TE exists which exhibits over 70% C_10_ free fatty acid in product distributions. While the truncated ClFatB3 exhibited a product distribution of only 50% of the ten-carbon species, activity-enhancing modifications could be coupled with subsequent mutagenesis efforts to tailor substrate specificity, ultimately yielding a highly active and highly specific decanoyl-ACP TE.

The N28S and I65M mutations identified in the active CupTE variant were incorporated into the truncated ClFatB3 sequence by cloning a D10S and I47M substitution into the expression vector (see **Figure S1**). Constructs ClFatB3_trunc_M1, ClFatB3_trunc_M2, and ClFatB3_trunc_M3 were generated to test the effect of D10S, I47M, and both D10S and I47M in combination, respectively. The effect of these mutations on the free fatty distribution is shown in **Figure 6**.

**Figure 6:**
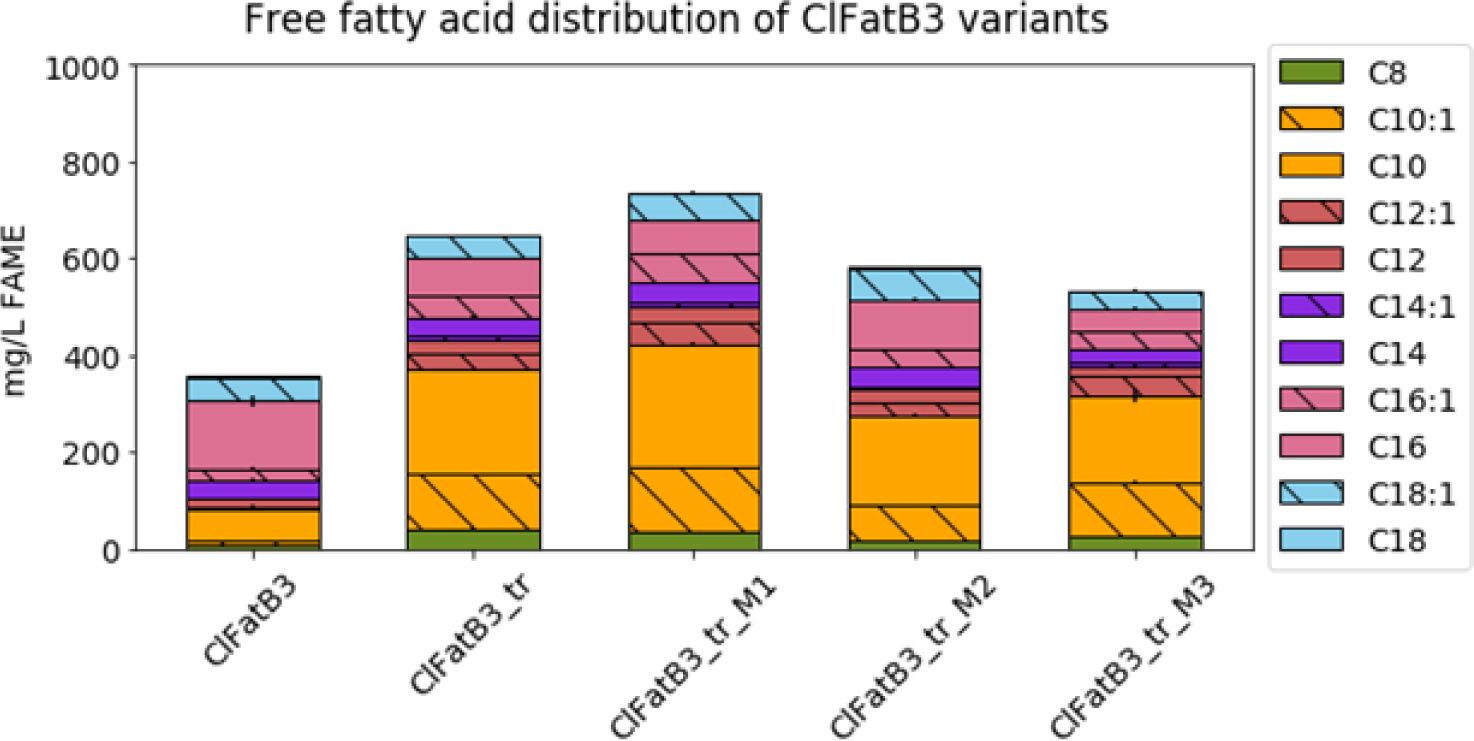
The effect of ClFatB3 variant on free fatty acid distribution in bulk cultures of RL08*ara E. coli* cells. Each TE was truncated at the location where the N-terminus aligned with the ΔA_54_ truncation in CupTE. Bar height represents the average titer obtained from biological triplicates, and error bars represent the standard error of the mean.

The D10S substitution on its own was shown to have the best effect in improving the overall titer in production cultures. This modification resulted in an 18% improvement in decanoic and decenoic acid titer compared to the truncated ClFatB3 variant and a 4.2-fold improvement when compared to the WT.

In summary, EnZymClass was used to successfully identify two unidentified medium-chain TE in the TE14 family in the ThYme database. These enzymes, ClFatB3 and CwFatB2, both originated from the *Cuphea* genus and demonstrated a substrate preference for decanoyl-ACP and dodecanoyl-ACP, respectively, when expressed in a fatty acid accumulating *E. coli* strain. Since both of these discoveries originated from a plant genus known to have medium-length acyl chains in its seed triglycerides, we wanted to ensure that EnZymClass was not simply returning sequences from *Cuphea* hosts and indeed was able to discern between medium and long-chain acyl-ACP TE homologs. To explore this, we tested ClFatB4, another TE from *Cuphea lanceolata* which was in the TE14 ThYme database family and therefore included in our model testing dataset. We also tested CpaFatB1, CpaFatB2A, and CpaFatB3, which are TEs from *Cuphea paucipetala* to sample other enzymes from the *Cuphea* genus. The free fatty acid distribution from ClFatB4, CpaFatB1, CpaFatB2A, and CpaFatB3, were all mostly dominated by the longer chemical species, thus confirming EnZymClass’ capability to classify substrate specificity among homologs from the same species **(Figure S2 and Table S4**).

To ratchet the activity of the novel TE discoveries made from EnZymClass’ predictions, the ClFatB3 and CwFatB2 genes were modified according to previous successful TE engineering efforts. The N-terminal truncation discovered from screening an octanoyl-ACP TE from *Cuphea palustris* for enhanced activity was thus incorporated into the predicted gene sequences. The truncated CwFatB2 variant displayed a favorable shift in product distribution toward dodecanoic acid. Furthermore, the truncated ClFatB3 variant resulted in a 3.3-fold improvement in C_10_ product titer over the WT. The final ClFatB3 variant, ClFatB3_trunc_M1, which included a D10S substitution also discovered in the CupTE screen by Hernández Lozada *et al.*, resulted in a 4.2- fold overall improvement in decanoic acid titer compared to the WT enzyme. Future work will entail the mutagenesis of ClFatB3 for enhanced substrate specificity to achieve near exclusive selectivity for C-_10_ FFA production in bacterial hosts.

## Discussion

With the rapid progress of sequencing technology, there have been a growing number of uncharacterized or unannotated protein sequences. Experimentally determining their functionality one at a time is both costly and time consuming. Computational methods that are capable of reliably predicting annotations from protein sequences are becoming increasingly indispensable. Machine Learning based supervised classification algorithms have evolved as the “go-to” method to solve this problem because of their ability to discern generalizable patterns from data without entrenched biases. These methods require a collection of characterized protein sequences from which they “learn” a set of rules or patterns employed to functionally annotate new uncharacterized sequences. However, the characterized datasets need to meet requirements such as class balance with enough instances per category of labels, feature set independence among others, to ensure generalizability of the ML model. Most common protein classification datasets that fail to meet at least one if not all of these specifications, lead to suboptimal performance of ML models on blinded test sets.

In this work, we presented a computational tool, termed EnZymClass, designed to address all the issues prevalent in protein classification pertinent datasets. To test the effectiveness of EnZymClass, we applied it on a specific protein classification task, categorization of plant acyl-ACP TEs into their respective substrate specificity groups. Expression of medium-chain specific acyl-ACP TE is the primary strategy for enabling development of biological platforms to high-value oleochemical derivatives. This approach has enabled enrichment of product distributions for medium-chain chemical species in bacteria, yeast, and plant systems (42, 64), (65). Several genes encoding for medium-chain TEs have been identified from plant species exhibiting medium-chain acyl chains in their seed oils (53), (66), (67). However, the vast majority of acyl-ACP TE in both prokaryotic and eukaryotic organisms exhibit long-chain specificity, preferring mostly 16 carbon substrates.

While the medium-chain activity is most desirable due to the relative scarcity of eight to twelve- carbon oleochemicals, most acyl-ACP TE exhibit a preference for C_14_ and above. Thus, several efforts have been made to explore genomes of plants with high fractions of the medium-chain oils to identify the TE gene responsible for the narrow substrate specificity (51), (42). While progress has been demonstrated in identifying the features which dictate specificity in acyl-ACP TE among plants (59), the throughput for bioprospecting, characterizing, and in some cases, engineering the acyl-ACP TE is largely inhibited by the testing pipeline, which requires derivatization of the free fatty acids into fatty acid methyl esters prior to analysis with gas chromatography (68). This prolonged testing cycle and the relative rarity of the medium-chain TE combined with the high value of medium-chain oleochemical derivatives has resulted in a broadly protected intellectual property landscape (51), (69), (70). Bioprospecting for novel medium-chain TE genes from organisms reported to have eight to ten carbon oil constituents is an alternative. While medium-chain-producing plants are well-catalogued (71), the bioprospecting process requires identification of an organism overlooked in the patent literature as well as the isolation of the desired gene from a cDNA library. The development of EnZymClass as a computational tool for identification of new medium-chain TE genes from primary sequence information simplifies the discovery process. When provided full-length amino acid sequences of putative acyl-ACP TE from the plant kingdom, EnZymClass can flag enzymes with the potential to produce distributions of over 50% of medium-chain free fatty acids in an *E. coli* production strain.

As part of this study, we compiled a characterized TE dataset consisting of 115 TE sequences and their corresponding substrate specificity category, long chain, medium chain or mixed. The assembled dataset is a fair representation of a typical protein classification dataset that exhibit attributes such as small size, class imbalance, high dimensionality, correlated features, which makes a conventional ML algorithm falter on unseen test sets. EnZymClass, on the other hand, displayed consistent performance when assessed through a rigorous validation strategy, devised so as to capture any trace of overfitting. We achieved classification accuracies (measured on the basis of correctly predicting the substrate specificity category of TEs) in the range of 0.52 to as high as 1 with mean accuracy of 0.80 and standard deviation of 0.06 across 10,000 simulations of our model validation scheme (discussed in **Methods**). EnZymClass also attained a mean precision score of 0.87 and mean recall score of 0.89 on medium-chain TE class prediction, the TE substrate specificity category that we are primarily interested in. In contrast an existing sequence similarity-based modeling approach achieved a mean accuracy, mean precision, and mean recall scores of 0.70, 0.79, and 0.79, respectively on medium-chain TE class prediction (detailed study given in **Results** section). We used EnZymClass to characterize all eukaryotic acyl-ACP TEs in the ThYme database (54). Among the three novel TEs predicted to be medium-chain specific by EnZymClass, two were experimentally validated to possess the desired activity.

Although we have attained reasonably high accuracy on TE substrate specificity classification task where the TEs were grouped into three different bins each representing a range of substrate specific chain lengths, we acknowledge that EnZymClass is currently unable to accomplish a deeper level of TE classification across specific chain lengths. It must be noted that the referred limitation to achieve a higher resolution of TE classification can be largely ascribed to a lack of characterized TE dataset with enough instances for each chain length. EnZymClass provides the flexibility to adapt to such requirements by simply modifying the base learners from classifiers to regressors. Moreover, the three categories of TE substrate specificity are not equally well predicted; prediction of the medium chained TE bin obtained the highest precision score of 0.87 followed by the long-chain category and the category of mixed specificity, which achieved mean precision scores of 0.78 and 0.54, respectively. Even though it can be viewed as a modeling limitation, the SVM base model hyperparameter “class weights” can be easily readjusted to impose further emphasis on the most poorly predicted category or the one that represents the subject of interest, thereby providing another technique to address class imbalance. In the TE substrate specificity prediction problem, EnZymClass was already biased towards the subject of our interest, medium-chain TEs. Hence, we decided against tuning the “class weight” hyperparameter. Furthermore, it has been previously recognized that primary sequence features alone may be insufficient to perfectly classify protein sequences and addition of protein structural features might boost prediction accuracy (72, 73). Taking this into account, a Convolutional Neural Network (CNN) base model (74) might be conveniently incorporated into the ensemble framework where the CNN extracts structural features from 3-D voxel representation of proteins, given that protein structural information is readily available.

EnZymClass can be adapted to other protein classification challenges ranging from general tasks such as protein structural class prediction or protein-protein interactions to more defined such as TE substrate specificity prediction or protein glycosylation site prediction (75). While general applications such as protein-protein interactions suffer from dataset imbalance (76), more specific tasks, for instance glycosylation site prediction may encounter yet another set of difficulties relating to small sized datasets. Issues related to high dimensionality and correlated feature set are ubiquitous in the protein classification domain. EnZymClass partially alleviates protein classification challenges while maintaining the computational efficiency required for swift functional characterization.

The identification of two previously uncharacterized medium-chain acyl-ACP TEs using only primary sequence information demonstrates the utility of ML in facilitating bioprospecting efforts, even when small training datasets of less than 200 datapoints are available. EnZymClass enabled the screening of 617 TE sequences *in silico* prior to experimental characterization of *in vivo* activity. With this work, we demonstrate how ML can be implemented to facilitate an enzyme engineering pipeline. The ML classification algorithm establishes suitable genes as templates for mutagenesis by identifying candidates which exhibit a desired function.

## Methods

### Dataset compilation

The training dataset included primary sequence and accompanying *in vivo E. coli* product distributions for 115 acyl-ACP plant TEs previously reported in scientific and patent literature (Table S6). Two of these TEs were previously tested for free fatty acid production prior to this study and were used in this study to supplement the training dataset. These include the sequences from *Auxenochlorella protothecoides* (KFM28838.1) and *Prunus sibirica L.* (AIX97815.1). The characterized dataset of 115 TEs is available at https://github.com/deeprob/ThioesteraseEnzymeSpecificity/tree/master/data. *E. coli* was chosen because it remains the most common and facile method for characterization of heterologous TEs. The product distribution data was subsequently used to classify each TE using discrete categories and a regression framework. For models which used discrete classification, the TEs were grouped into one of three categories. The “medium-chain” category contained TE which resulted in distributions of at least 50% C_8_ to C_12_ free fatty acids. The “long-chain” category contained TE which produced 50% C_14_ to C_18_ free fatty acids and less than 10% C_8_ to C_12_ free fatty acids. Finally, the “mixed distribution” category contained TE which yielded distributions between 10% and 50% C_8_ to C_12_ free fatty acids. In the implementation of the linear regression classifier, each sequence was assigned a number which represented the fraction of the total free fatty acid distribution constituted of C_8_ to C_12_ free fatty acids.

### Feature extraction

In this work, 47 alignment-free feature extraction techniques that encode primary sequence information of the enzymes into fixed-length feature vectors were employed. The feature extraction techniques fall under four categories, Kernel methods, N-gram methods, Physicochemical encoding methods, and PSSM profile based methods. TE sequence feature encoding was conducted utilizing source codes from three open-source python or R-based tools, KeBABS (77), iFeature (78), and POSSUM (79). While KeBABS is already an existing R package for Kernel methods, we have developed three PyPI (80) packages (ifeatpro (81), ngrampro (82) and pssmpro (83)), one for each of the three other categories that encompasses 44 additional feature extraction techniques apart from the three Kernel based methods used in EnZymClass, enabling straightforward numerical encoding of protein sequences. Information about accessibility and usage of the three protein sequence-encoding packages is provided in **Supplementary Information**. The feature extraction category, name, software package used to deploy them and literature from which they are adopted are listed in **Table 2**. A brief description of the 47 feature extraction techniques divided into their respective categories is provided in **Text S1**.

**Table 2:** 47 feature extraction techniques were used in EnZymClass. The feature extraction type, name, software used to create it, and literature from which it was adopted is provided in this table.

| Type | Name | Software Used | Literature Reference |
| --- | --- | --- | --- |
| Kernel | Spectrum Kernel | KeBABS | (14) |
| Kernel | Mismatch Kernel | KeBABS | (19) |
| Kernel | Gappy Pair Kernel | KeBABS | (84) |
| N-gram | Kmer | ngrampro | (26) |
| N-gram | GAA-kmer | ngrampro | NA |
| Physicochemical | AAC | ifeatpro | (85) |
| Physicochemical | CKSAAP | ifeatpro | (86) |
| Physicochemical | TPC | ifeatpro | (85) |
| Physicochemical | DPC | ifeatpro | (87) |
| Physicochemical | DDE | ifeatpro | (87) |
| Physicochemical | GAAC | ifeatpro | (88) |
| Physicochemical | CKSAAGP | ifeatpro | (78) |
| Physicochemical | GTPC | ifeatpro | (78) |
| Physicochemical | GDPC | ifeatpro | (78) |
| Physicochemical | Moran | ifeatpro | (89) |
| Physicochemical | Geary | ifeatpro | (90) |
| Physicochemical | NMBroto | ifeatpro | (91) |
| Physicochemical | CTDC | ifeatpro | (92–94) |
| Physicochemical | CTDT | ifeatpro | (92–94) |
| Physicochemical | CTDD | ifeatpro | (92–94) |
| Physicochemical | CTriad | ifeatpro | (95) |
| Physicochemical | KSCTriad | ifeatpro | (78) |
| Physicochemical | SOCNumber | ifeatpro | (78) |
| Physicochemical | QSOrder | ifeatpro | (78) |
| Physicochemical | PAAC | ifeatpro | (96) |
| Physicochemical | APAAC | ifeatpro | (96) |
| PSSM-based | AAC-PSSM | pssmpro | (97) |
| PSSM-based | DPC-PSSM | pssmpro | (97) |
| PSSM-based | AADP-PSSM | pssmpro | (97) |
| PSSM-based | PSSM-AC | pssmpro | (98) |
| PSSM-based | PSSM-CC | pssmpro | (99) |
| PSSM-based | RPSSM | pssmpro | (100) |
| PSSM-based | Tri-gram | pssmpro | (101) |
| PSSM-based | EDP | pssmpro | (102) |
| PSSM-based | EEDP | pssmpro | (102) |
| PSSM-based | MEDP | pssmpro | (102) |
| PSSM-based | TPC-PSSM | pssmpro | (103) |
| PSSM-based | AATP | pssmpro | (103) |
| PSSM-based | k-separated bigrams | pssmpro | (104) |
| PSSM-based | D-FPSSM | pssmpro | (105) |
| PSSM-based | S-FPSSM | pssmpro | (105) |
| PSSM-based | PSE-PSSM | pssmpro | (106) |
| PSSM-based | DP-PSSM | pssmpro | (107) |
| PSSM-based | PSSM-Composition | pssmpro | (108) |
| PSSM-based | Smoothed-PSSM | pssmpro | (109) |
| PSSM-based | AB-PSSM | pssmpro | (110) |
| PSSM-based | RPM-PSSM | pssmpro | (110) |

### EnZymClass: <u>E</u>nsemble method for <u>e</u>nZyme <u>Class</u>ification

#### Model description

EnZymClass is comprised of *N* base learners which provide their outputs to a meta learner that predicts the functional attribute of proteins (enzyme specificity class in our case). Although all base learners are trained using the same principle (SVM or NN or GBT), the heterogeneity among them is governed by the different feature extraction techniques (described in the **Feature Extraction** section) used to encode the set of protein sequences. Each feature extraction technique creates a unique feature vector representation of proteins which serve as input to a designated base learner in EnZymClass. The base learner trained on the set of encoded protein sequences yields the predicted functional attribute of a given protein sequence as an output. The outputs of the *k*- best base learners are passed on to the meta learner that uses a majority voting scheme (described in **The Meta Learner** section) to predict protein functional attribute category. The *k*-best base learners are selected on the basis of their performance on validation set as described in **Model Training** section. EnZymClass pipeline is presented in **Figure 7**.

**Figure 7:**
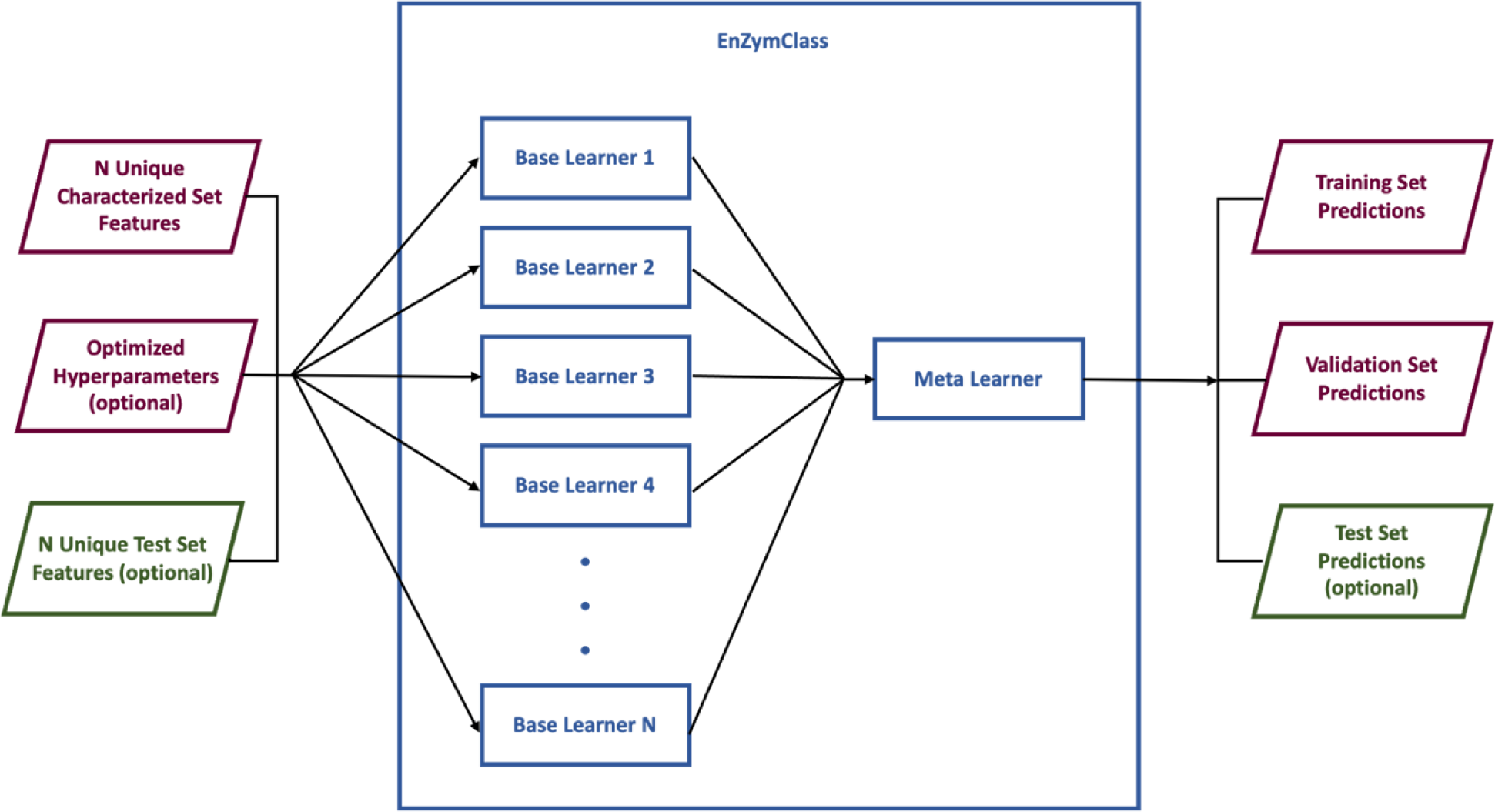
EnZymClass pipeline is described here. Unique feature representation technique designated Base Learners get independently trained and predict functional attributes of protein sequences. The independent predictions are collected by the Meta Learner that outputs the final model prediction by calculating the majority vote of the predictions made by each individual Base Learner. EnZymClass requires a labelled characterized set of sequences as input. Users can also provide an unlabeled test set and optimized hyperparameters for each Base Learner as optional arguments. Training and validation sets are created from subsets of the characterized “training” set. EnZymClass outputs model predictions for the training set, validation set and test set (if provided).

#### Base learner algorithms

1. Support Vector Machine: The Support-Vector-Machine-based learner of functional attribute prediction included Principal Component Analysis (PCA) for dimensionality reduction of the feature space followed by a Support Vector Classifier (111) to predict protein class. The PCA based dimensionality reduction step was carried out to decrease the number of parameters required to train a SVM model and make EnZymClass more generalizable. The one-versus-one strategy was used to adapt the SVM for multi-class classification (112). The number of PCA components, SVM model kernel, regularization parameter C, and kernel coefficient gamma were selected by optimizing these hyperparameters using a 5-fold cross validation scheme (described in **Model Training** section).
2. Neural Network: The Neural Network-based learner of functional attribute prediction included Principal Component Analysis for dimensionality reduction of the feature space followed by an Artificial Neural Network classifier to predict protein class. The PCA-based dimensionality reduction step was carried out to decrease the number of parameters required to train a NN model and make EnZymClass more generalizable. The number of PCA components, hidden layer size of NN, initial learning rate, and L2 regularization parameter alpha were selected by optimizing these hyperparameters using a 5-fold cross-validation scheme (described in **Model Training** section).
3. Gradient Boosting Trees: The Gradient-Boosting-Tree-based learner of functional attribute prediction included Principal Component Analysis for dimensionality reduction of the feature space followed by a Gradient Boosting Tree classifier to predict protein class. The PCA based dimensionality reduction step was carried out to decrease the number of parameters required to train a NN model and make EnZymClass more generalizable. The number of PCA components, number of estimators or decision trees to consider, learning rate, and maximum depth of the trees were selected by optimizing these hyperparameters using a 5-fold cross-validation scheme (described in **Model Training** section).

#### The Meta learner

The meta learner accepts the outputs of all the base learners as an input vector, implements a hard majority voting scheme on the input vector, and returns the consensus prediction of the label (TE substrate specificity in our case) as an output. The meta learner output is calculated as follows:

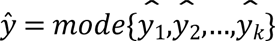

Where *ŷ* is the output of the meta learner, *ŷ*_*i*_ is the output of the base learner *i*, *k* is the number of best performing base learners to consider, and *mode* is a function that selects the most frequent value among a set of values.

#### Model hyperparameters

EnZymClass has several hyperparameters which can be tuned to improve model performance. It has a hyperparameter *k* that denotes the number of best performing base learners that the meta learner needs to take into account. Each base learner has a number of hyperparameters depending on the learning algorithm used to train them. The respective learning algorithm-dependent hyperparameters of the base learners are shown in **Table 3**. The hyperparameters are learnt through a 5-fold cross validation scheme discussed in **Model Training** section.

**Table 3:**
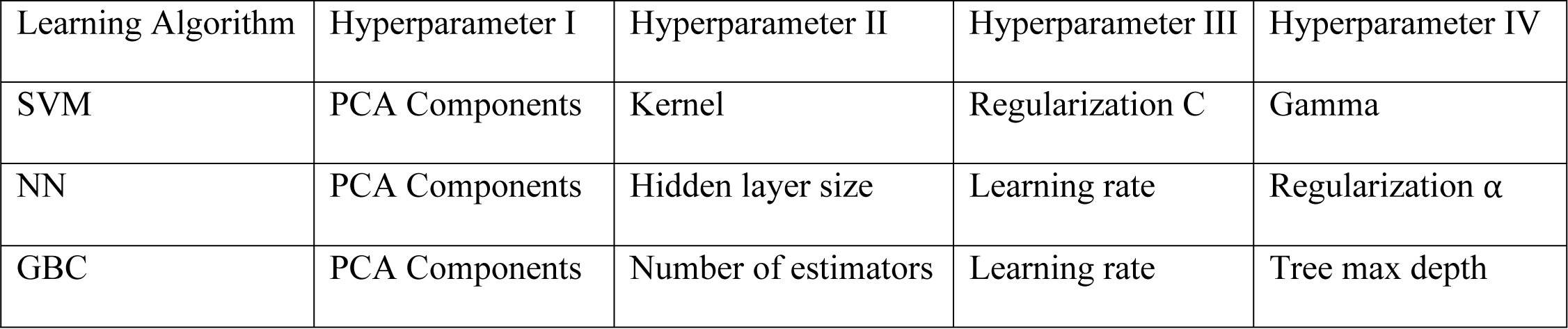
The hyperparameters of each base learner are dependent on the learning algorithm used to train it. The learning algorithm-dependent hyperparameters are displayed here.

#### Model evaluation metrics

The performance of EnZymClass was measured using four popular classification metrics, 1) accuracy score, 2) precision score on the medium chain TE class, 3) recall score on the medium chain TE class, and 4) Matthew’s correlation coefficient (MCC).

Accuracy score for a multi-class classification problem is defined as:

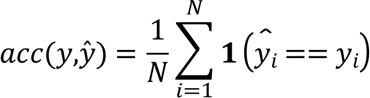

Where *ŷ*_*i*_ is the predicted value for sample *i*, *y*_*i*_ is the corresponding true sample, *N* is the number of samples, and **1** is an indicator function that is equals to 1 if a certain condition is true and 0 otherwise.

Precision score for a class *i* is defined as:

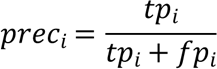

Where *tp*_*i*_ is the number of true positives for the *i*-th class and *fp*_*i*_ is the number of false positives for the *i*-th class.

Recall score for a class I is defined as:

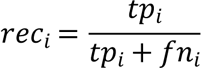

Where *fn*_*i*_ is the number of false negatives recorded for the *i*-th class.

Matthew’s Correlation Coefficient for a multiclass classification problem is defined as:

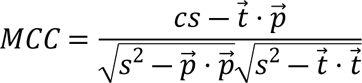

Where, *c* = ∑_*k*_ *c*_*kk*_ or the total number of instances correctly predicted across all *k* classes, *s* = ∑_*i*_ ∑*c*_*i,j*_ or the total number of instances, *t*_*k*_ = ∑_*i*_ *c*_*ik*_ or the number of times class *k* actually occurred, *p*_*k*_ = ∑_*i*_ *c*_*ki*_ or the number of times class *k* was predicted, and *C* is a confusion matrix of size *k* x *k*.

#### Model training

EnZymClass can be trained using python’s numpy and scikit-learn modules (113, 114). Model training and validation can be divided into five stages described below.

1. Random seed assignment and dataset division: EnZymClass requires a characterized dataset as input. It divides the characterized dataset (115 TE enzyme sequences labeled according to their corresponding substrate specificity category in this study) into training and validation set by a 75-25 percentage split. A random seed can be specified in EnZymClass to reproduce results. Changing the random seed will produce different training and validation sets, an event which will be used later to evaluate model performance.
2. Feature representation: The training and validation set of sequences needs to be encoded by different feature representation techniques before feeding them as input to EnZymClass. In the current work, we used feature representation methods described in the **Feature Extraction** section to transform TE sequences into 47 distinct feature vector representation. The distinct feature vectors of the training set of sequences were used to train 47 separate base learners operating on the same principle (PCA+SVM or PCA+NN or PCA+GBT).
3. Base model training: The base models accept the feature vector representation of protein sequences as input and predicts their functional attributes. They are trained using scikit-learn dedicated modules for PCA, SVC, NN and GBC. Each base model is a scikit-learn pipeline object consisting of PCA instance followed by the learning algorithm instance (SVC, NN, or GBC). The pipeline object was trained on the feature vector representation of the training set of TE sequences in this study. EnZymClass allows the hyperparameters of the base learners to be optimized using the GridSearchCV module of scikit-learn with a 5-fold cross-validation process.
4. Meta learner prediction: The trained base models in EnZymClass independently predict the functional attributes of protein sequences and pass on their predictions to the Meta Learner that uses a hard-voting based majority vote classifier to output the final prediction of the functional attribute. In the present study, the 47 base models trained only on the training set were used to independently predict the substrate specificity category of enzymes in both training and validation sets. The output predictions of these base learners were passed on to the meta learner. The parameter *k* of EnZymClass, representing the *k*-best base models to pass on to the meta learner, was chosen to be 5.
5. Model evaluation: EnZymClass allows model evaluation by providing methods to calculate four popular classification metrics, accuracy score, precision score, recall score, and MCC. In the current study, the validation set accuracy, precision score (on the medium-chain TEs) and recall score (on the medium-chain TEs) of the 47 base learners and the ensemble model were recorded. The entire Model Training procedure was repeated 10,000 times by varying the random seed specified initially. This resulted in different training and validation sets, thus affecting model performance and yielding a distribution of training and validation set accuracies, precision, and recall scores for 47 base learners and EnZymClass. The objective of evaluating our model multiple times by varying the training and validation set was to check its robustness to the training set. A parametric sweep of the ensemble model parameter *k* was performed to illustrate its effect on validation score.

#### Model prediction

To predict uncharacterized protein sequences in a test set, the test sequences need to be converted to feature vectors using the same feature extraction techniques employed to encode sequences in the training set. The *N* base models delegated to each feature vector representation are subsequently trained and hyperparameter optimized using all the characterized protein sequences. The trained base models are used to independently predict the functional attributes of test set proteins. The output predictions of the *k* best-performing base learners are passed on to a meta learner that uses a hard-voting based majority vote classifier to output the final prediction of the functional attribute. In the present work, we followed the above procedure to obtain substrate specificity predictions of uncharacterized TE sequences. The parameter *k* was selected based on the results of the parametric sweep study discussed in the **Model Training** section.

### Computational study workflow

In the current study, we employed EnZymClass to predict substrate specificity of plant acyl-ACP TEs by training it on a small, unbalanced TE dataset with high sequence similarity between the enzyme sequences. EnZymClass is specifically built to tackle the challenges of high-dimensionality, small-sized and imbalanced datasets, correlated features, and high sequence similarity among proteins at every stage of its pipeline. We used 47 alignment-free feature extraction techniques, proven to be effective in multiple application areas of protein sequence classification, to numerically encode TE sequences such that EnZymClass can effectively extract as much information from primary sequences as realistically possible. To prevent overfitting, the feature vectors generated through the extraction process were decomposed into lower dimensional and linearly uncorrelated features using Principal Component Analysis. The lower dimensional and decomposed set of feature vectors were used to train individual base learners and independently predict substrate specificity of TEs. An ensemble method was used to circumvent the dataset imbalance problem and retain high prediction accuracy. The generalizability of EnZymClass was assessed through a rigorous model validation strategy (discussed in **Model Training** section) where we simulated 10,000 different versions of training and validation datasets from the characterized set of TE sequences. We recorded the performance of the framework on these validation datasets using three popular classification metrics, accuracy, precision, and recall. The entire workflow of the study including model training, validation, and prediction of uncharacterized TE sequences is demonstrated in **Figure 8**.

**Figure 8:**
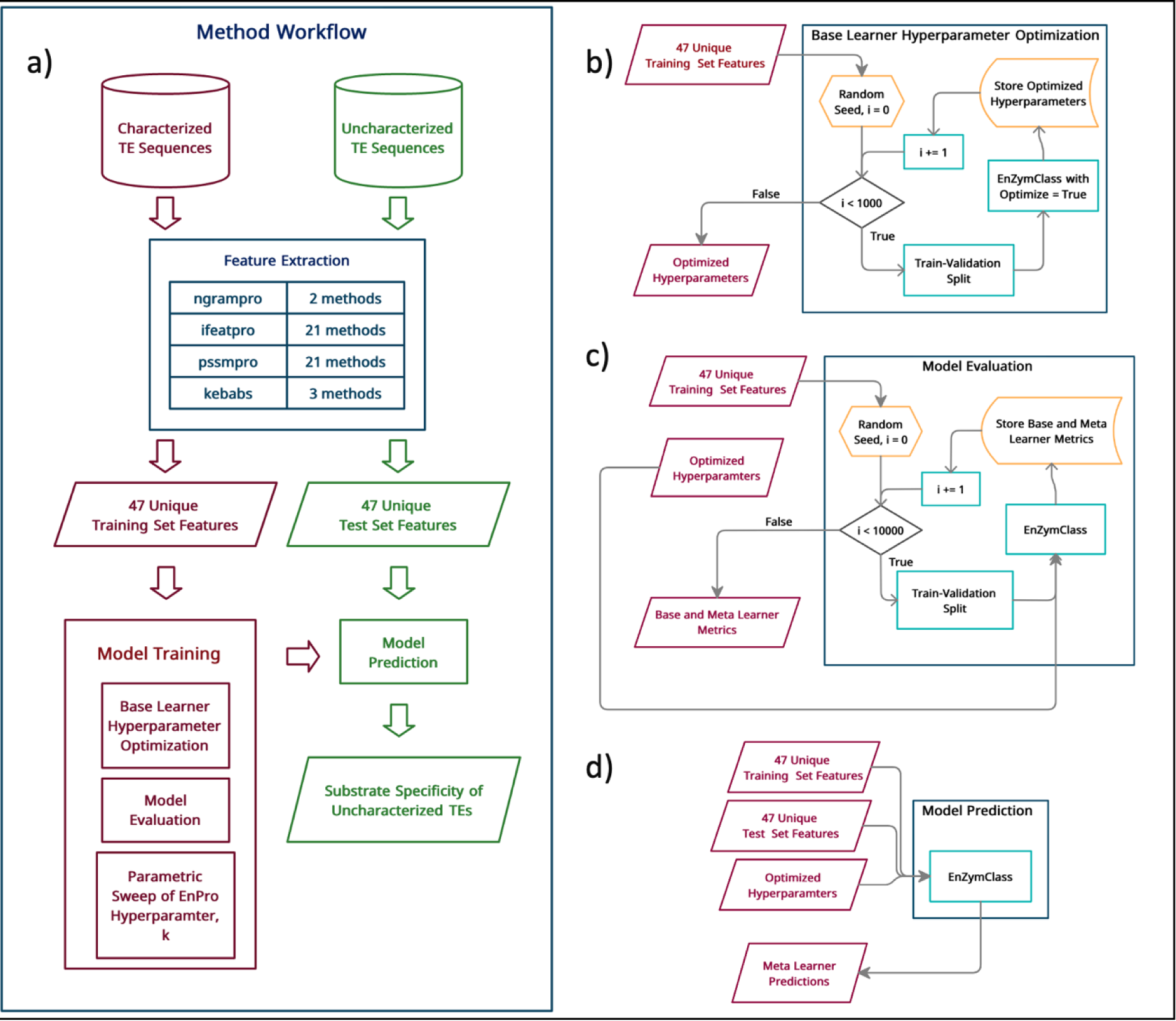
a) outlines the entire workflow of TE substrate specificity prediction method. Both characterized and uncharacterized TE sequences are encoded into 47 unique training and test feature sets. In the Model Training phase, the training feature sets are used to optimize hyperparameters of EnZymClass in addition to evaluating model predictions on a validation set created by splitting the original training set into two segments. A parametric sweep of the EnZymClass hyperparameter k, the number of best base learners that passes on their predictions to the meta learner, is also performed. Finally, the optimized hyperparameters along with the entire training set is used to train EnZymClass and predict substrate specificity of uncharacterized TE sequences in the Model Prediction phase; b) describes the Base Learner Hyperparameter Optimization component of the Model Training phase. The unique set of features are divided into different training and validation sets, 1000 times. Each time, EnZymClass is trained, and the base learners are hyperparameter optimized using a training set. The optimized hyperparameters are stored for each run. At the end of 1000 runs, the most frequent hyperparameters among the list of hyperparameters for each base learner are selected as the chosen hyperparameters; c) illustrates the Model Evaluation component of the Model Training phase. The unique set of features are divided into different training and validation sets, 10000 times. Each time, EnZymClass is trained using the training set and the optimized hyperparameters obtained in the Base Learner Hyperparameter Optimization stage. The resulting model predictions of the Base and Meta Learners in EnZymClass are noted and they are used to calculate classification metrics that indicate model performance. The list of metrics for each run is stored and a distribution of classification metrics are returned after 10,000 runs; d) depicts the Model Prediction phase. The entire Training feature set along with the previously optimized hyperparameters are used to train EnZymClass which predicts the substrate specificity of the test set enzymes.

EnZymClass’ novel pipeline and model architecture allows it to overcome challenges posed by small, characterized protein sequence datasets for the purpose of attaining high prediction accuracies. It employs 47 alignment-free descriptors of protein sequences, the maximum number of encoders used by any protein classification algorithm known, with the ability to automatically select the encoders best suited for a specific protein classification task. Intuitively, the ability of EnZymClass to select the best set of feature descriptors should allow it to generalize across protein classification datasets covering a wide range of application areas in computational biology. It is worth noting that although deep learning algorithms such as CNNs or RNNs can automatically build features descriptive of a specific classification task, they require characterized datasets ranging from thousands to millions of instances. unlike EnZymClass which can work with as few as a hundred training instances as illustrated in this study. Moreover, unlike similar ensemble learning algorithms described in (28,115–119), EnZymClass includes an in-built decomposition technique to tackle issues related to high-dimensionality and is capable of self-tuning model hyperparameters, a phenomenon that often leads to better and more robust model performance. Additionally, it provides the user with the flexibility of choosing the base learner training algorithm among SVM, GBC and NN.

### Media and molecular biology materials

Media components were ordered from IBI Scientific (Dubuque, IA) and Fisher Scientific (Waltham, MA). Oligonucleotides were purchased from Integrated DNA Technologies (Coralville, IA) and gene fragments were purchased from Twist Bioscience (San Francisco, CA). Enzymes for DNA cloning were purchased from New England Biolabs (Ipswich, MA).

### DNA Cloning and *in vivo* characterization of Acyl-ACP TEs

All TE homologs were tested in the high copy pTRC99a expression vector (120). Gene fragments were cloned via restriction digestion with *HindIII* and *EcoRI* enzymes and ligated into the pTRC99a construct in place of the BTE gene. Point mutations and truncations were incorporated into the ClFatB3, CwFatB2, and BTE genes by designing mutagenic primers which yielded PCR amplicons amenable to Gibson Assembly (121).

Once the sequence was verified, each TE construct was subsequently transformed into RL08*ara*, an *E. coli* MG1655 derivative deficient in the acyl-CoA synthetase gene to enable accumulation of free fatty acids (122). Single colonies of the RL08 transformants were grown overnight at 37°C and 250 r.p.m. in 5 mL of LB media supplemented with 100 mg/L of carbenicillin. All strains and vectors used in this study are in **Table S3**.

To begin free fatty acid production trials, shake flasks containing 25 mL of LB supplemented with 4 g/L of glycerol were inoculated with 275 µL of the stationary phase culture. The cultures were then allowed to grow at 30°C and 250 r.p.m. until they reached an OD of 0.2 – 0.3. Isopropyl β-D-thiogalactoside IPTG was then added to the media to a final concentration of 20 µM to induce transcription of the TE genes. After an additional 24 hours of culturing at 30°C and 250 r.p.m., the flasks were removed, and 2.5 mL of the media was sampled for derivatization and characterization of the free fatty acid distribution. For analysis of the free fatty acid distribution present in the supernatant, 10 mL of sample were centrifuged at 4500 x *g* for 20 minutes, and 2.5 mL of the clarified supernatant was collected. The fatty acid quantification method described by Politz *et al.* was then followed with slight modifications (68). Namely, the internal standard solution of odd-chain free fatty acids was prepared by combining heptanoic, nonanoic, undecanoic, tridecanoic acid to a final concentration of 5 g/L in methanol, and pentadecanoic and heptadecanoic acid were added to a final concentration of 1 g/L. 100 µL of the internal standard was added to each 2.5 mL sample. After the GC-FID data was obtained, each even-chain free fatty acid was quantified by normalizing the peak area of its derivatized methyl ester to the adjacent peaks of the odd-chain methyl esters. For example, decanoic acid concentration was quantified by first obtaining the concentration to peak area ratio of the undecanoic acid and tridecanoic acid methyl esters. The peak area of the dodecanoic acid methyl ester peak was then multiplied by the averaged concentration to peak area ratio of both the C11 and C13 peaks to calculate the concentration of dodecanoic acid present in the media. To ensure accuracy of the method, recovery standards were added to 2.5 mL of fresh media each time the above protocol was performed. The recovery standard solution consisting of 5 g/ L of octanoic, dodecanoic, dodecenoic, and tetradecanoic acid and 1 g/L of decanoic and hexadecenoic acid. 100 µL of the recovery standard and 100 µL of the internal standard solutions were added to the 2.5 mL blank media samples. These recovery samples were then processed alongside the samples from the shake flask cultures.

## Supporting Information captions

**Text S1**: Description of the 47 feature extraction techniques used in EnZymClass

**Table S1**: A list of three performance metrics for the 47 base models ranked by their mean accuracy score on varying validation datasets

**Table S2:** Performance of EnZymClass trained on different base learners.

**Table S3**: A list of *E*. coli strains and expression plasmids used in this study.

**Table S4**: Fatty acid methyl ester distributions and titers obtained from derivatization of bulk stationary phase culture following *in vivo* culture experiments

**Table S5**: Fatty acid methyl ester distributions and titers obtained from derivatization of culture supernatants following *in vivo* culture experiments.

**Table S6:** Fatty acid methyl ester distributions and titers obtained from various *in vivo* culture experiments used to train EnZymClass.

**Figure S1**: Multiple sequence alignment of BTE, ClFatB3, ClFatB3-2, and CwFatB2 to CpFatB1 generated by Clustal Omega

**Figure S2**: The effect of various TE homologs from the Cuphea genus on free fatty acid distribution in bulk cultures of RL08ara E. coli cells.

## Acknowledgments

The sequences for *Auxenochlorella protothecoides* (KFM28838.1) and *Prunus sibirica L.* (AIX97815.1) were synthesized by the U.S. Department of Energy Joint Genome Institute (JGI), a DOE Office of Science User Facility, supported by the Office of Science of the U.S. Department of Energy. Computations for this research were performed on the Pennsylvania State University’s Institute for Computational and Data Sciences’ Roar supercomputer.

## Author contributions

DB and MAJ conceived the study, analyzed the data, and wrote the manuscript. MAJ mined the literature for the training dataset. DB mined the ThYme database for the test dataset, designed and simulated the computational model and workflow. MAJ designed and ran the experimental procedures. AJL provided guidance during the initial conceptualization and implemented a preliminary model to validate the concept. BFP and CDM obtained the funding, provided valuable insights and wrote the manuscript. DB and MAJ contributed equally to this work.

